# Heterogeneity in viral infections increases the rate of deleterious mutation accumulation

**DOI:** 10.1101/2021.05.07.443113

**Authors:** Brent Allman, Katia Koelle, Daniel Weissman

## Abstract

RNA viruses have high mutation rates, with the majority of mutations being deleterious. We examine patterns of deleterious mutation accumulation over multiple rounds of viral replication, with a focus on how cellular coinfection and heterogeneity in viral output affect these patterns. Specifically, using agent-based intercellular simulations we find, in agreement with previous studies, that coinfection of cells by viruses relaxes the strength of purifying selection, and thereby increases the rate of deleterious mutation accumulation. We further find that cellular heterogeneity in viral output exacerbates the rate of deleterious mutation accumulation, regardless of whether this heterogeneity in viral output is stochastic or is due to variation in cellular multiplicity of infection. These results highlight the need to consider the unique life histories of viruses and their population structure to better understand observed patterns of viral evolution.

## 2 Introduction

RNA viruses have high mutation rates and undergo frequent population bottlenecks, making them particularly prone to the accumulation of deleterious mutations. As such, these populations can experience deleterious mutation loads, which is the burden on fitness that recurrent and persistent mutations have on populations (Crow, 1958; Agrawal and Whitlock, 2012) Indeed, the accumulation of deleterious mutations in viruses has been repeatedly demonstrated using experimental evolution. In particular, experiments have demonstrated that serial population bottlenecks impact rates of deleterious mutation accumulation in viral populations (Chao, 1990; Clarke et al., 1993; Escarmís et al., 1996; Elena et al., 1998; Poon and Chao, 2004; García-Arriaza et al., 2005). Drugs that exploit this accumulation by increasing already high mutation rates can drive viral populations extinct (Anderson et al., 2004; Pauly and Lauring, 2015; Bank et al., 2016). Experimental studies have also shown that cellular coinfection affects the rate of deleterious mutation accumulation in viral populations (Wilke and Novella, 2003; Novella et al., 2004). In particular, cellular coinfection leads to slower purging of deleterious mutations because selection is relaxed: when multiple viral genomes are present in a cell, they all share their protein products (Zavada, 1976; Froissart et al., 2004). With multiple copies of the same gene that have differential fitness, phenotypes and genotypes of the offspring will not necessarily be matched. Cellular coinfection therefore allows for “phenotypic hiding” of deleterious mutations (Wilke and Novella, 2003; Novella et al., 2004).

Several processes reduce the accumulation of deleterious mutations in RNA viruses. One such mechanism is through the evolution of higher fidelity polymerase proteins, thus reducing deleterious mutation rates (Pfeiffer and Kirkegaard, 2003; Coffey et al., 2011; Cheung et al., 2014). Recombination (and its segmented analogue, reassortment) also reduces the rate of deleterious mutation accumulation through the generation of high fitness viral genotypes via viral sex. Bylimiting cellular multiplicity of infection (MOI), superinfection exclusion (Turner et al., 1999; Schaller et al., 2007; Folimonova, 2012)also reduces the opportunity for phenotypic hiding. However, superinfection exclusion also limits the opportunity for viral sex to occur, and thus its net effect on the rate of deleterious mutation accumulation is unknown.

The effect of cellular MOI on the rate of deleterious mutation accumulation is particularly interesting to consider given its uniqueness to viral populations and that cellular coinfection is, in effect, a double-edged sword: providing an opportunity for viral sex to occur, while increasing the extent of phenotypic hiding. Here, we develop a model to examine the effects of cellular coinfection on deleterious mutation accumulation in viral populations in the context of these opposing effects. We first show that the simplest version of the model recapitulates previous findings in the literature (Wilke and Novella, 2003; Novella et al., 2004), that indicate that cellular coinfection, in the absence of genetic exchange, increases the accumulation of deleterious mutations. We then extend this model to include cellular heterogeneity in viral output, based on recent experimental findings that demonstrate extreme cellular heterogeneity in response to viral infection (Russell et al., 2018; Martin et al., 2020). We find that heterogeneity, whether due to variation in cellular MOI or intrinsic cellular variation, increases the rate of deleterious mutation accumulation. Our findings highlight how life history characteristics common to viruses can impact deleterious mutation accumulation.

## 3 Model

### 3.1 Base Model

We use a generalized Wright-Fisher model of the viral population (Fig. 1), with *V* virions infecting a host cell population of size *C*. Both *V* and *C* remain constant over time, yielding a constant average multiplicity of infection (MOI) of *V/C*. Each virion has *g* genes in its genome. These genes are distributed across *y* freely reassorting gene segments, with no recombination within segments. Deleterious mutations occur at a rate of *U/g* per gene per generation, such that the overall deleterious mutation rate occurs at a rate of *U* per genome per generation. In simulations of this model, we use *y* ∈ {1, 2, 4, 8} to capture a range of reassortment potentials, with *y* = 8 reflective of influenza A virus genomes. For simplicity, we use *g* = 8 in all simulations so that genes can be evenly distributed across the considered segments. Within each gene, we adopt an infinite sites assumption. Thus each genome can be characterized simply by how many deleterious mutations it carries at each of its *g* genes.

**Figure 1:**
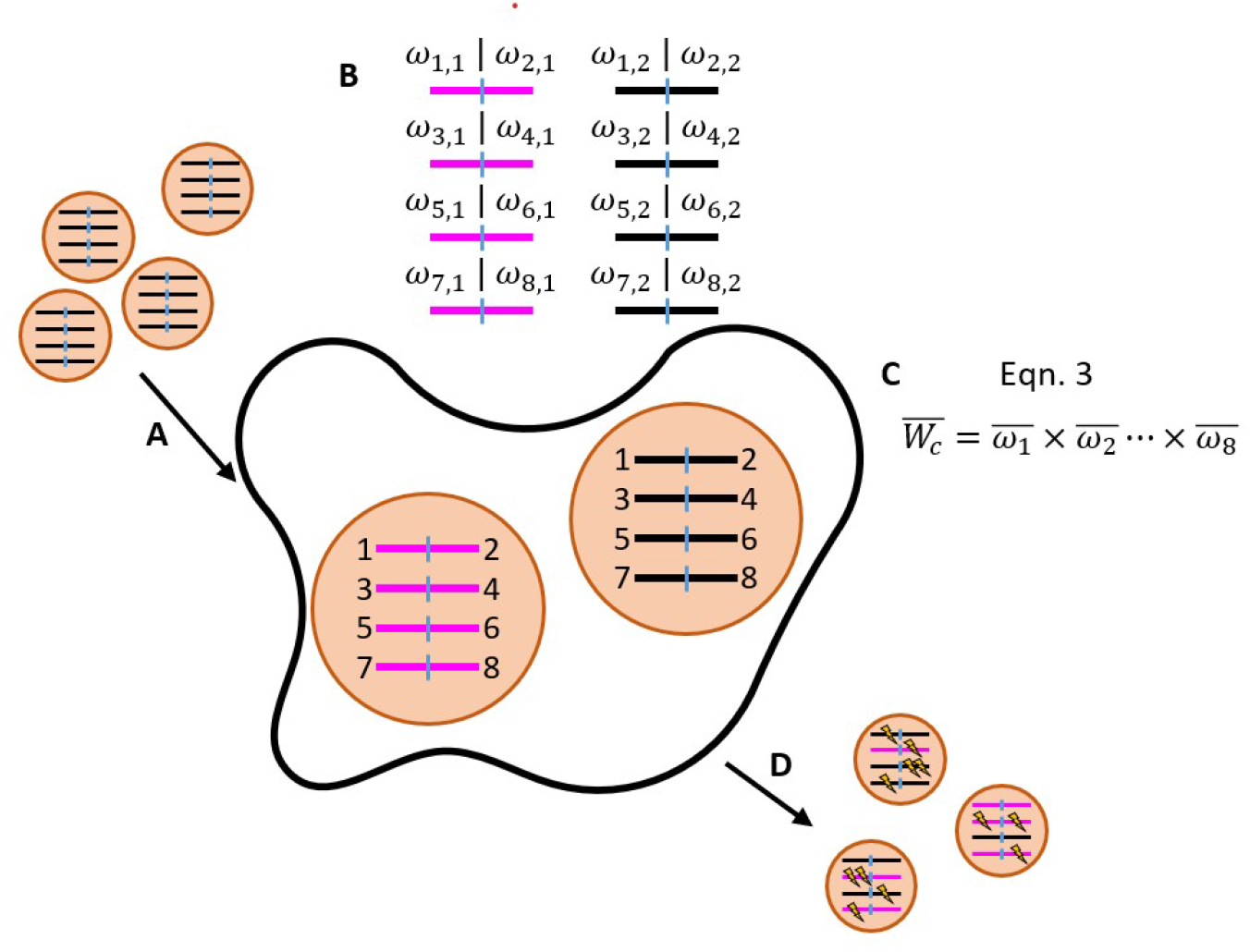
Schematic of the base model with a viral genome over a single generation. Each generation consists of a series of steps A-D. (A) *V* virions infect *C* cells. Here, two virions infect the shown cell. The viral genomes each have *g* = 8 genes distributed across *y* = 4 gene segments. Each gene is labeled 1-8. (B) Within each cell, the fitnesses of individual gene copies are calculated using equation (1). These *ω*_*i,j*_ values will be used to calculate the group fitness for each gene. (C) For each gene in each cell, average fitnesses are calculated using equation (2). Cellular fitnesses are then calculated using equation (3). (D) *V* viral progeny are formed by selecting parental cells according to their cellular fitnesses, and then selecting gene segments at random from within the cell. Deleterious mutations (lightning bolts) are introduced during the formation of these viral progeny. Steps A-D are repeated for *t* generations.

At the beginning of each generation, the *V* virions are randomly assigned to the *C* cells, resulting in a Poisson distribution of virions across cells. Once inside the cells, the numbers of mutations on each gene determine the aggregate fitness of the viral population within each cell. This aggregate fitness, which we call “cellular fitness”, determines the relative contribution of each cell’s virus population to the next generation of virions. To calculate cellular fitness, we first calculate the fitness of each gene that was delivered to a cell:

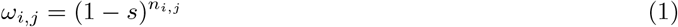

where *s* is the constant fitness cost of a deleterious mutation and *n*_*i,j*_ is the number of deleterious mutations on gene *i* delivered by virion *j*. For each gene *i*, we calculate the mean fitness of the gene in a cell as

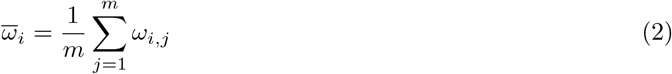

where *m* is the multiplicity of infection of the host cell. Finally, we calculate the expected cellular fitness, *W*_*c*_, as:

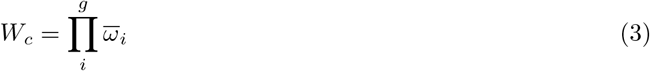

Equations 1 - 3 make three key assumptions: (1) each mutation within a gene contributes multiplicatively to the fitness of the cell (Eqn. 1), (2) each copy of a gene *i* contributes equally to 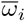 via incomplete dominance (Eqn. 2), and (3) each gene segment is essential and equally important in its contribution to cellular fitness (Eqn. 3). We make these assumptions based on the idea that when multiple virions of differing genotypes infect a cell, the produced viral proteins are treated as common goods used in the generation of progeny virions.

At the end of each generation, we draw the *V* progeny virions for the next generation from across the set of infected cells. Each progeny virion is drawn independently, with the probability that the virion comes from cell *c* proportional to *W*_*c*_. Given that the virion comes from cell *c*, each of its *y* gene segments is drawn randomly from the parental virions that infected the cell. As such, a high fitness gene segment is as likely to be drawn from a cell as a low fitness gene segment, reflecting our assumption that cellular fitness depends on the aggregate of shared viral proteins that have been produced in a cell. Once all parental gene segments have been chosen, the mutations are added as described above. We repeat this full process for *t* discrete generations.

### 3.2 Heterogeneous Cellular Output Stemming from Differences in Cellular Characteristics

Viral output from cells can be affected by host cell characteristics such as size, cell type, and cell cycle stage (Brooke et al., 2013; Schulte and Andino, 2014; Heldt et al., 2015; Golumbeanu et al., 2018; Leviyang and Griva, 2018; Russell et al., 2018; Xin et al., 2018; Phipps et al., 2019; Vera et al., 2019). To consider the effect of heterogeneity in virus output on deleterious mutation accumulation, we extend our base model described above by adapting an approach used by Lloyd-Smith et al. (2005) to describe population-level viral transmission heterogeneity (superspreading dynamics). Specifically, we introduce cellular heterogeneity by making a distinction between the *cellular output* 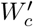 and the cellular fitness *W*_*c*_. For each cell *c*, the cellular fitness *W*_*c*_ is still determined by the genes of the infecting viruses according to Eqn. 3 as above. But in the next generation, the probability that a viral progeny is drawn from *c* is instead proportional to 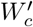, a gamma-distributed random variable with mean *W*_*c*_ and shape parameter *k*, i.e., probability density function:

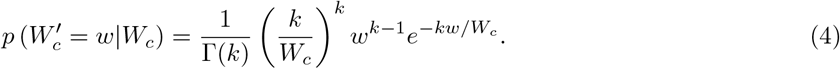

The parameter *k* controls the extent of cellular heterogeneity. As *k* → ∞, heterogeneity driven by host cell characteristics becomes minimal and probability that a viral progeny derives from cell *c* converges to the fitness, 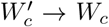. In contrast, as *k* → 0, the probability that a viral progeny derived from cell *c* becomes increasingly dependent on host cell characteristics and relatively less dependent on the genotypes of viral genes delivered to a cell.

### 3.3 Heterogeneity in Cellular Output Stemming from Differences in Cellular Multiplicity of Infection

Virus output from cells can also be affected by cellular multiplicity of infection, with higher cellular MOI having the potential to increase viral yield (Phipps et al., 2019; Martin et al., 2020). To consider the effect that this source of cellular heterogeneity in virus output may have on deleterious mutation accumulation, we extended the base model to allow cellular multiplicity of infection to impact cellular output. Specifically, we let cellular output of a cell with multiplicity of infection *m*_*c*_ be given by a linear relationship between cellular input and cellular output, 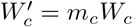. While numerous other functional forms are possible, this is the simplest one that allows us to assess the qualitative effect of input-dependence on deleterious mutation accumulation.

## 4 Results

In our results, we focus on the mean number of deleterious mutations accumulated in a viral population by generation *t*. Unless specified otherwise, data shown are from the final generation of the simulated infection, *t* = 20 or *t* = 150. With a viral generation being approximately 5 hours long for viruses such as influenza (Baccam et al., 2006), this corresponds to approximately 4 days post-infection and 31 days post-infection, respectively. In addition to *t* = 20 and *t* = 150 conveniently approximating the number of generations over acute and chronic infections, we choose these two endpoints due to substantial changes in rates of deleterious mutation accumulation over time. Roughly, *t* = 20 is the time to approach mutation-selection balance for many of our simulations, so changes in the number of accumulated mutations at this time reflect shifts in mutation-selection balance. At the later time *t* = 150, we can distinguish between populations with a slow-acting Muller’s ratchet versus ones with a fast-acting Muller’s ratchet.

### 4.1 Phenotypic hiding relaxes selection

We first show that our base model reproduces key findings on deleterious mutation accumulation from previous work using similar cellular coinfection modeling frameworks, in addition to classical population genetics. That is, we establish that the sizes of the virus and host cell populations influence the rate of genetic drift and the extent of phenotypic hiding in the context of cellular coinfection.

For simplicity, we begin by considering an unsegmented genome (*y* = 1), so there is no reassortment. One key finding from the field of population genetics is that reducing population size increases the rate of deleterious mutation accumulation due to increased genetic drift, particularly in asexual populations (Fisher, 1930; Wright, 1931; Kimura et al., 1963; Lynch et al., 1995). This effect has mostly been studied under purely individual-level selection. This is a good approximation of our system at low MOI, where most infected cells are infected by only a single virion. Indeed simulations of our model reproduce this effect of population size at low MOI (Fig. 2A).

**Figure 2:**
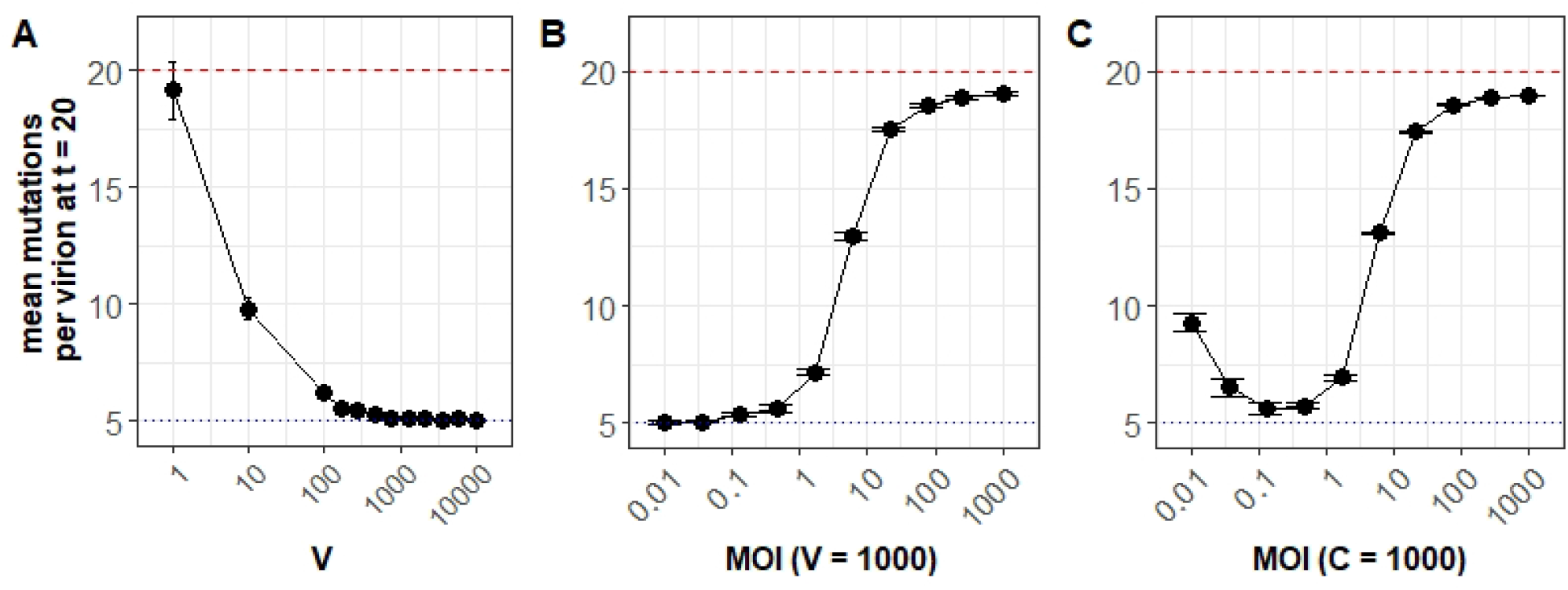
Simulated patterns of deleterious mutation accumulation without cellular heterogeneity. **(A)-(C)** Mean number of accumulated mutations at *t* = 20 generations. Each data point shown is the average across 20 replicate simulations with error bars showing the standard error, except for the three largest population sizes in subplot (C), which have only a single replicate shown due to computational limitations. Red dashed lines show the theoretical expectation of mutation accumulation at selective neutrality (*Ut*). Blue dotted lines show the expectation of mutation accumulation for an infinite viral population size at its mutation-selection balance (*U/s*). Parameter values are *V* = 1000, *C* = 1000, *U* = 1, *s* = 0.2, *g* = 8, *y* = 1 unless otherwise indicated. **(A)** MOI (= *V* /*C*) is kept constant at 0.1, such that cell population sizes scale linearly with viral population sizes. Higher viral population sizes have lower rates of deleterious mutation accumulation. Large viral populations reach their deterministic mutation-selection balance and have lower rates of deleterious mutation accumulation thereafter. **(B)** The virus population size is kept constant at *V* = 1000 and cell population size *C* is modified to change MOI. Here, increasing MOI increases phenotypic hiding and therefore deleterious mutation accumulation. **(C)**. The cell population size is kept constant at *C* = 1000 and the virus population size *V* is modified to change MOI. Here there is a tradeoff at low MOI, genetic drift, whose sole effects are shown in (A), dominates and mutation accumulation rates are high because of small viral population sizes. At high MOI, phenotypic hiding, whose effects are shown in (B), dominates and mutation accumulation rates are high because of high levels of cellular coinfection.

Previous work has shown that cellular coinfection and the sharing of viral proteins relaxes the strength of selection on individual virions, and thus allows deleterious mutations to accumulate at a faster rate in viral populations than otherwise expected (Wilke and Novella, 2003; Froissart et al., 2004; Novella et al., 2004). Our model recapitulates this “phenotypic hiding” in simulations where the viral population size is kept constant and the number of cells is modified to change the overall MOI (Fig. 2B). The monotonic increase in the number of accumulated deleterious mutations in the population observed with increases in MOI is directly attributable to relaxed selection.

In Figure 2A, we found that increases in the viral population size (while maintaining a constant MOI) can slow mutation accumulation by decreasing the rate of genetic drift and slowing deleterious mutation accumulation. In Figure 2B, however, we found that increases in MOI (while maintaining a constant viral population size) can accelerate mutation accumulation by increasing the extent of phenotypic hiding. Thus, increases in viral population size that are not matched by increases in the size of the cell population could yield a non-monotonic relationship between viral population size and the rate of deleterious mutation accumulation. Figure 2C shows the results of this tension between the effects of genetic drift and phenotypic hiding. At low MOI (≪1), coinfection is rare, and the primary effect of an increase in the viral population size is a reduction in the strength of genetic drift, decreasing mutation accumulation. As MOI approaches 1, however, phenotypic hiding starts to play a more pronounced role and mutation accumulation increases. At very high MOI (≳ 1) phenotypic hiding is essentially complete and deleterious mutations accumulate at the neutral rate *U*.

The rate of deleterious mutation accumulation should decrease in segmented viral genomes because reassortment can re-create high fitness genotypes that have been lost to drift by combining segments that have a small number of deleterious mutations, halting Muller’s ratchet (Chao, 1990). We confirm that this occurs in our base model when we consider the viral genome of *g* = 8 genes divided across *y* = 1, 2, 4, 8 gene segments (Fig. 3). Because reassortment does not affect the approach to mutation-selection balance, it has little effect at early times (e.g., *t* = 20), but at later times it results in a slower ‘clicking’ of the ratchet. As such, more highly segmented genomes, which allow more reassortment, have lower levels of accumulated deleterious mutations than genomes that have fewer gene segments.

**Figure 3:**
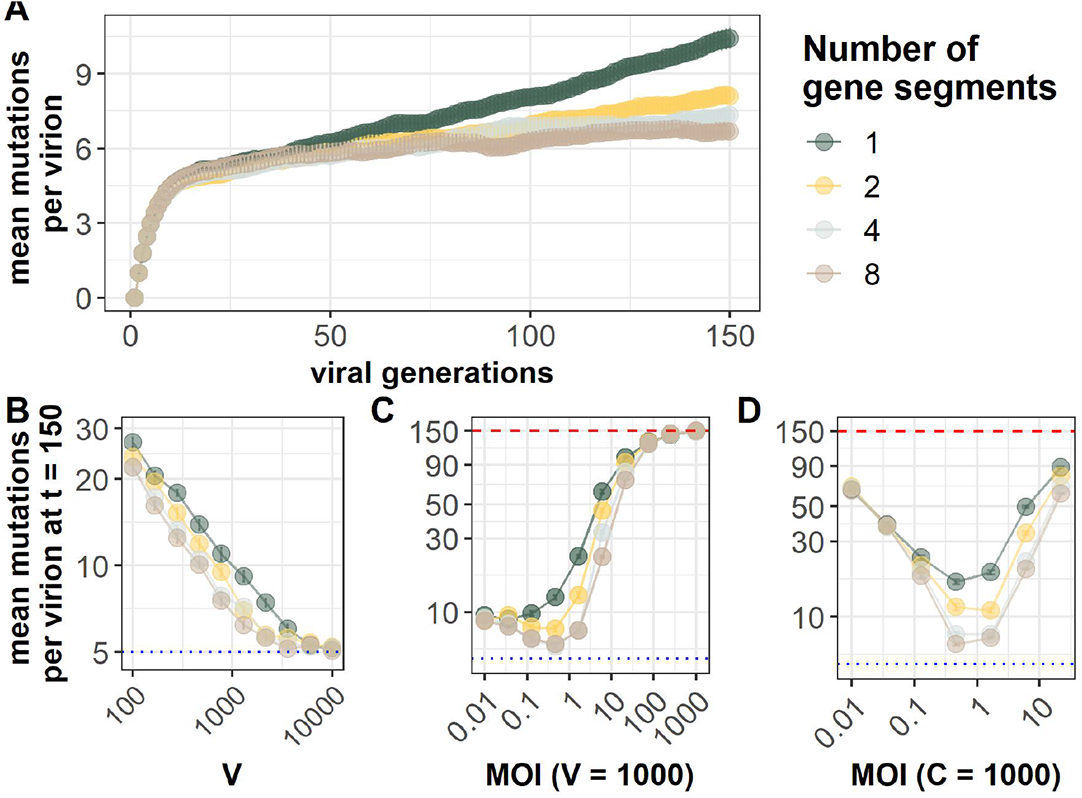
Genome segmentation slows the accumulation of deleterious mutations. In (A)–(D), the per genome mutation rate is *U* = 1 and the fitness cost of mutations is *s* = 0.2. Each data point is the average across 20 replicate simulations with error bars showing the standard error. Red dashed lines in (C) and (D) show the theoretical expectation of mutation accumulation at selective neutrality (*Ut*). Blue dashed lines in (B) - (D) show the expectation of mutation accumulation for an infinite viral population size at its mutation-selection balance (*U/s*). **(A)** Average number of deleterious mutations accumulated over time at a viral population size *V* = 1000 and a cell population size of *C* = 10000 for varying numbers of segments. **(B)–(D)** show the average number of deleterious mutations harbored by a viral population at generation *t* = 150 under different parameters. **(B)** Reassortment slows mutation accumulation in small populations subject to Muller’s ratchet. MOI (= *V* /*C*) is kept constant at 0.1 by scaling linearly the cell population size *C* proportionally with the viral population size *V*. **(C)–(D)** Mutation accumulation is slowest at intermediate MOI *≈* 0.3, balancing the effects of reassortment and phenotypic hiding. In **(C)**, MOI is varied by changing *C*, while in **(D)** it is varied by changing *V*. At high MOI *≳* 1, phenotypic hiding is nearly complete and mutations accumulate at close to the neutral rate.

Reassortment has the largest effect on mutation accumulation at intermediate viral population sizes that are large enough to effectively select against individual mutations but small enough to be vulnerable to Muller’s ratchet, 1*/s < V < e*^*U/s*^*/s* (Barton and Otto, 2005). At larger viral population sizes, the ratchet clicks very slowly even in non-reassorting viruses, and therefore reassortment provides little benefit (Fig. 3B) (Muller, 1964). The effect is observable even when cellular coinfection is rare, consistent with findings from the population genetic literature (Bell, 1988; Charlesworth et al., 1993; Cohen et al., 2006).

Our model also recapitulates the impact of sex that are described by classical population genetics (Fisher, 1930; Wright, 1931; Kimura et al., 1963; Lynch et al., 1995). The capacity for viruses to have sex influences deleterious mutation accumulation as MOI increases. In Figure 3C, we keep the viral population size the same as we test different sized cell populations to modulate MOI. To this end, as coinfection events become more common at moderate MOI, segmented genomes accumulate fewer deleterious mutations than their unsegmented counterparts. These findings are supported by previous work which shows that sex (here, reassortment) slows rates of deleterious mutation accumulation and Muller’s ratchet by reseeding the least-loaded classes of individuals with deleterious mutation loads (Muller, 1964; Chao et al., 1997).

However, segmented genomes are still vulnerable to the impacts of phenotypic hiding (Fig. 3C). While genomes that are segmented accumulate fewer deleterious mutations than unsegmented populations at high MOI, the segmented genomes do not continue to gain a benefit from increased coinfection (Fig. 3C, D). It remains true that segmented populations can slow the accumulation of deleterious mutations by allowing the least-loaded gene segments to unlink from their less fit background. This reseeds more fit classes of individuals, but their ability to be maintained is still impacted by the strength of selection. The same amount of phenotypic hiding is occurring in the segmented populations at high MOI, so selection approaches a neutral limit as in unsegmented populations (Fig. 2C, D). Sex provides no benefit to mutation accumulation when mutations are selectively neutral.

### 4.2 Stochastic heterogeneity increases deleterious mutation accumulation

As described in the Model section above, we consider the effect of heterogeneity driven by host cell characteristic by integrating individual cell heterogeneity with virus-driven differences in cellular fitness using draws from a gamma distribution, parameterized with dispersion parameter *k*.

As expected, simulations with *k* ≳ 1 behave like the ones described above that do not incorporate stochastic heterogeneity (Fig. 4). However, for *k* ≪1, more deleterious mutations accumulate under all tested conditions (Fig. 4A and Supplementary Figure 1). This is because the increased stochasticity reduces the efficacy of purifying selection. This has little effect under very high mean MOI ≳ 1, because phenotypic hiding already weakens selection such that mutations accumulate at nearly the neutral rate, but it can greatly increase mutation accumulation at lower MOI where selection would otherwise be strong enough to halt mutation accumulation.

**Figure 4:**
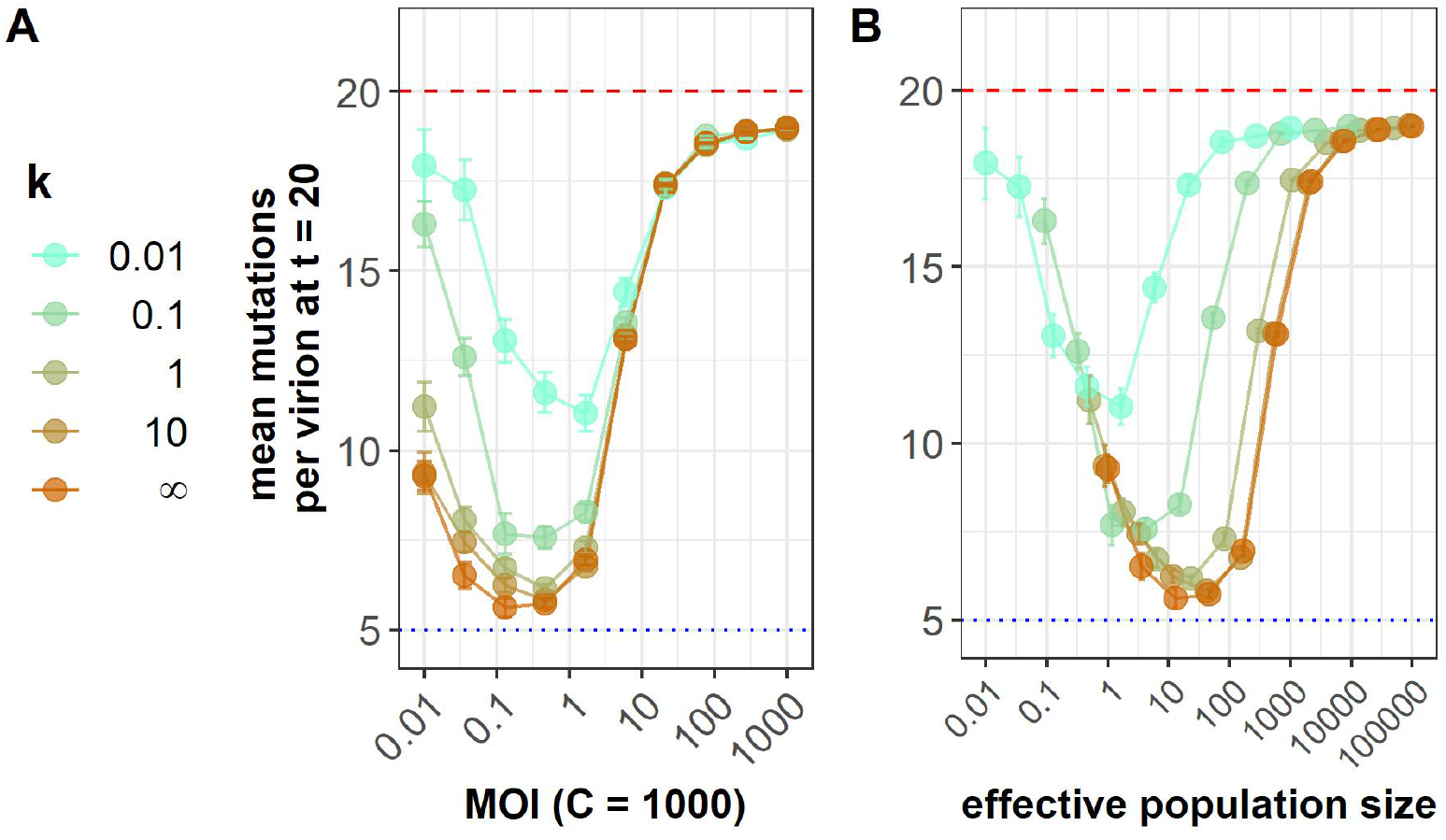
Stochastic heterogeneity increases deleterious mutation accumulation. All panels show mean number of deleterious mutations after *t* = 20 generations of within-host infection. Stochastic heterogeneity is parametrized by *k*, with *k ≪*1 corresponding to strong heterogeneity and *k* = *∞* corresponding to the base model without heterogeneity. **(A)** Stochastic heterogeneity has the largest effect at low MOI. At high MOI, phenotypic hiding makes selection ineffective even in the absence of heterogeneity. **(B)** Same data as (A), but shown as a function of the predicted effective viral population size *V*_*e*_ = *V/*(1 + 1*/k*). The collapse of the different curves on the left side of the plot shows *V*_*e*_ accurately captures the effect of heterogeneity on mutation accumulation in the regime where the effect is strong. In both panels, each data point shown is the average of 20 replicate simulations with error bars showing the standard deviation (with the exception of the three largest population sizes in which show only a single simulation). Red dashed lines show the theoretical expected mutation accumulation at selective neutrality (*Ut*). Blue dashed lines show the average number of mutations for an infinite viral population size at its mutation-selection balance (*U/s*). Parameters are *C* = 1000, *U* = 1, *s* = 0.2, *y* = 1.

The effect of stochastic cellular heterogeneity on mutation accumulation can be better understood by quantifying the effective viral population size, *V*_*e*_ in these simulations. Stochastic heterogeneity in cellular virus production increases the variance in offspring number among virions *σ*^2^, and thereby decreases the viral effective population size, given by *V*_*e*_ ≡ *V/σ*^2^, where *σ*^2^ is more generally the variance in the offspring distribution (Ewens, 1982). We can calculate *σ*^2^ at low MOI (≪1) where almost all cells are infected with either zero or one virion. At low MOI and ignoring fitness differences between virions, the offspring distribution is a gamma-Poisson mixture (i.e., a negative binomial) with a mean of 1 (because *V* is constant) and variance *σ*^2^ = 1 + 1*/k* since each infected cell produces a gamma-distributed number of virions (each cell with a different mean) and the virions infect an approximately Poisson-distributed number of cells in the next generation.

We tested different population sizes under a constant cellular population size of *C* = 1000 and computed *V*_*e*_. This allows us to see how phenotypic hiding is contributing to a reduction in effective population size in the context of stochasticity. In Figure 4B, we conclude that stochastic heterogeneity drives differences in mutation accumulation primarily at low and intermediate MOI, when viral population sizes are small and little to no coinfection takes place. This intuitively makes sense because we know that small populations are susceptible to stochastic effects like drift (Supplementary Figure 1). Importantly, despite increases in stochasticity that reduce effective population sizes at high MOI, phenotypic hiding results in similar numbers of mutations accumulating in the high MOI regimes. At high MOI, phenotypic hiding puts populations into a regime of selective neutrality despite their large realized population sizes (Supplementary Figure 1). We know that phenotypic hiding is driving this effect because different values of our dispersion parameter reduce the effective population sizes, but at high MOI similar numbers of mutations still accumulate (Fig. 4A and B). So regimes with both high population size and high MOI show us that stochasticity can reduce effective population sizes, but phenotypic hiding is primarily responsible for deleterious mutation accumulation.

One may notice that the small populations can at times produce *V*_*e*_ *<* 1. This is because for very small *k* and small population sizes, we no longer follow a negative binomial offspring distribution; so our estimation of *V*_*e*_ breaks down for these small populations with small *k*. However, this does not impact the results in Figure 4 which shows that stochastic heterogeneity increases rates of deleterious mutation accumulation at low and intermediate viral population sizes.

### 4.3 Input-dependent viral populations accumulate slightly more mutations at intermediate MOI

We next performed simulations under cellular heterogeneity that stems from differences in viral input. When we assumed that viral output scaled linearly with viral input, we found slightly more mutations accumulated compared to the base model at intermediate MOI (Fig. 5 and Supplementary Figure 2). In contrast, at both high and low MOIs, there were no appreciable differences observed between the number of mutations observed in the base model relative to those in the input-dependent model considered here.. These results can be understood as follows.

**Figure 5:**
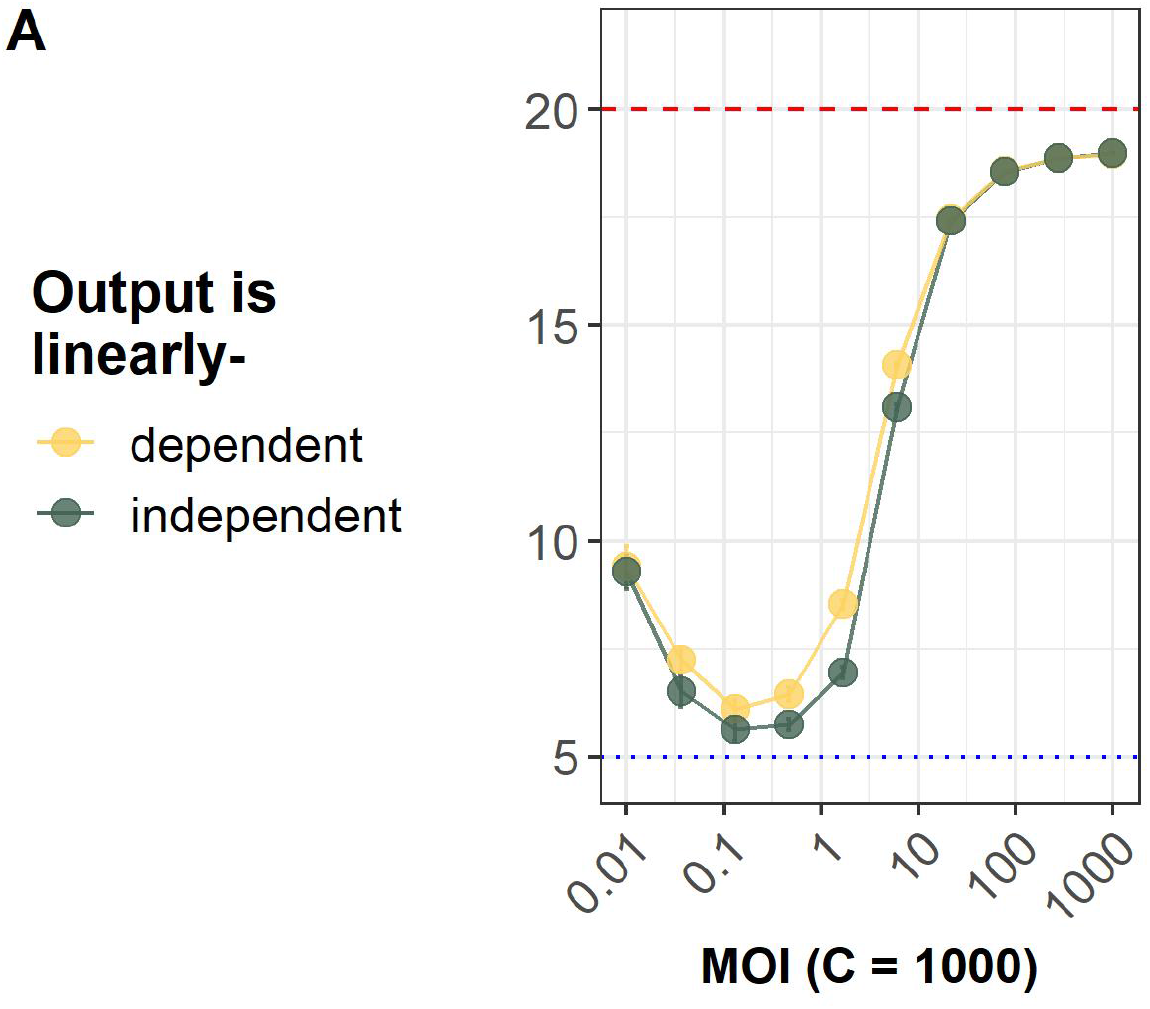
Input-dependent cellular fitness values increase the rate of deleterious mutation accumulation slightly at intermediate MOI. The black line show results from the base model. The yellow line shows simulations where cellular fitness *W*_*c*_^*l*^ depends both on the amount of viral input *m*_*c*_ to the host cell and the average fitness of the viral genomes in the host cell *W*_*c*_. We use the per genome mutation rate *U* = 1 and the fitness cost of mutations *s* = 0.2. Each data point shown is the average across 20 replicate simulations with error bars showing the standard error. Red dashed lines show the theoretical expectation of mutation accumulation at selective neutrality (*Ut*). Blue dashed lines show the expectation of mutation accumulation for an infinite viral population size at its mutation-selection balance (*U/s*). **(A)** Average number of mutations harbored by an individual virion at generation *t* = 20. The cell population size is kept constant at *C* = 1000 and virus population sizes are modified to change MOI.

At low MOI (≪1), almost all infected cells are infected by only a single virion, so the input-output relationship is irrelevant (Supplementary Figure 2A). At very high MOI (≳ 1), phenotypic hiding is nearly complete such that mutations accumulate near the neutral rate in both models (Supplementary Figure 2B). At intermediate MOI, however, there is a mix of singly infected cells, where virions do not experience phenotypic hiding, and multiply infected cells, where virions experience phenotypic hiding. With viral output scaling linearly with viral input, the multiply infected cells contribute more viral progeny to the next generation, thereby contributing viral genomes that have experienced relaxed selection

## 5 Discussion

Here we consider how cellular coinfection in viral infections impacts deleterious mutation accumulation using an *in silico* simulation model. Using our model, we were able to recapitulate previous results of relaxed selection that occurs under regimes of phenotypic hiding (Wilke and Novella, 2003; Froissart et al., 2004; Novella et al., 2004). We then extended these findings by showing that the heterogeneity inherent to viral infections, including cellular heterogeneity and differences in production of virions due to variation in number of infecting viral particles, increases the rates of deleterious mutation accumulation during viral infections.

Segmentation and reassortment reduce selection interference among genes by allowing more fit variants to jointly reproduce progeny that do not contain all of the deleterious mutations harbored by their parents Turner (2003). However, our simulations indicate that phenotypic hiding can drastically reduce this benefit of segmented genomes (Fig. 3C). We show that intermediate levels of coinfection (MOI ≈ 0.3) are optimal for segmented viral populations since they allow sex to occur frequently enough to reduce interference without significant levels of phenotypic hiding.

While our extensions to the base model have incorporated some realism, our model remains highly simplified. In particular, there are two key features of natural infections which impact population dynamics that we have still not incorporated. First, many infections show substantial spatial structure (reviewed in Gallagher et al. (2018)). This could result in high MOI hotspots, increasing the potential for both phenotypic hiding and, in segmented viruses, reassortment. However, spatial structure also means that coinfecting virions are likely to be close relatives, reducing both the negative impact of phenotypic hiding and the benefits of reassortment.

The second key aspect of natural infections not captured by our model is that we assume a constant viral population size, while natural infections expand from a small inoculum. Population expansion has been shown to increase the number of segregating deleterious mutations in the population but also decreases the per individual number of deleterious mutations (Gazave et al., 2013). We do not know how population expansion would interact with cellular coinfection. Interpreting our results over a dynamic range of MOI using Figure 2C indicates that if a viral population were to grow in a limited cell population, selection would first be at an individual-level in a small population susceptible to drift, and then selection could weaken over the course of an infection as viral population size increases. On the other hand, it is unclear what would happen if the viral population were to continue to colonize new tissue as it grew such that MOI remained roughly constant.

One possible genetic extension of our model would be to include epistasis among mutations. Positive epistasis would result in additional mutations accumulating because the fitness effect of adding a new mutation decreases with each subsequent mutation. Negative epistasis would have the opposite effect: selection would be more strict and thus fewer mutations would accumulate.

Phenotypic hiding can be seen as an example of social interactions between viruses at the intracellular level. The emerging field of “sociovirology” examines how such interactions between viruses, including during cellular coinfection, can have an impact on the evolution of viral populations (Vignuzzi et al., 2006; Andino and Domingo, 2015; Bordería et al., 2015; Díaz-Muñoz et al., 2017; Sanjuán, 2017; Aguilera and Pfeiffer, 2019). The importance of coinfection in viral evolution has been demonstrated empirically (Chao et al., 1997; Turner et al., 1999; Wilke and Novella, 2003; Froissart et al., 2004). Specifically, cellular MOI depends on viral traits such as aggregation via collective infectious units (reviewed in Sanjuán (2017)), while other traits may limit coinfection via superinfection exclusion (Sun and Brooke, 2018). Some of the other modern work in the field also highlights the role of heterogeneity (Andreu-Moreno and Sanjuán, 2018; Sun and Brooke, 2018). However, while much of sociovirology focuses on positive selection, our work shows that interactions among virions also have large effects on the ability of purifying selection to shape the evolution of viral populations.

## Data Availability

The code used to produce the data shown in this paper was written and implemented in MATLAB R2020a and is available at https://github.com/allmanbrent/coinfection_heterogeneity. Visualization was performed using R version 4.0.1.

## Acknowledgments

BEA thank members of the Koelle lab (particularly Molly Gallagher and Jeremy Harris) for helpful comments on implementation of the model.

## Funding

BEA was supported by the NSF National Science Foundation Graduate Research Fellowship Program (grant no. DGE-1444932). DBW was supported by a Simons Investigator Award in the Mathematical Modeling of Living Systems.

## Conflicts of Interest

The authors report no conflicts of interest.

## Supplementary Materials

### 1 Stochastic Heterogeneity

**Figure 1:**
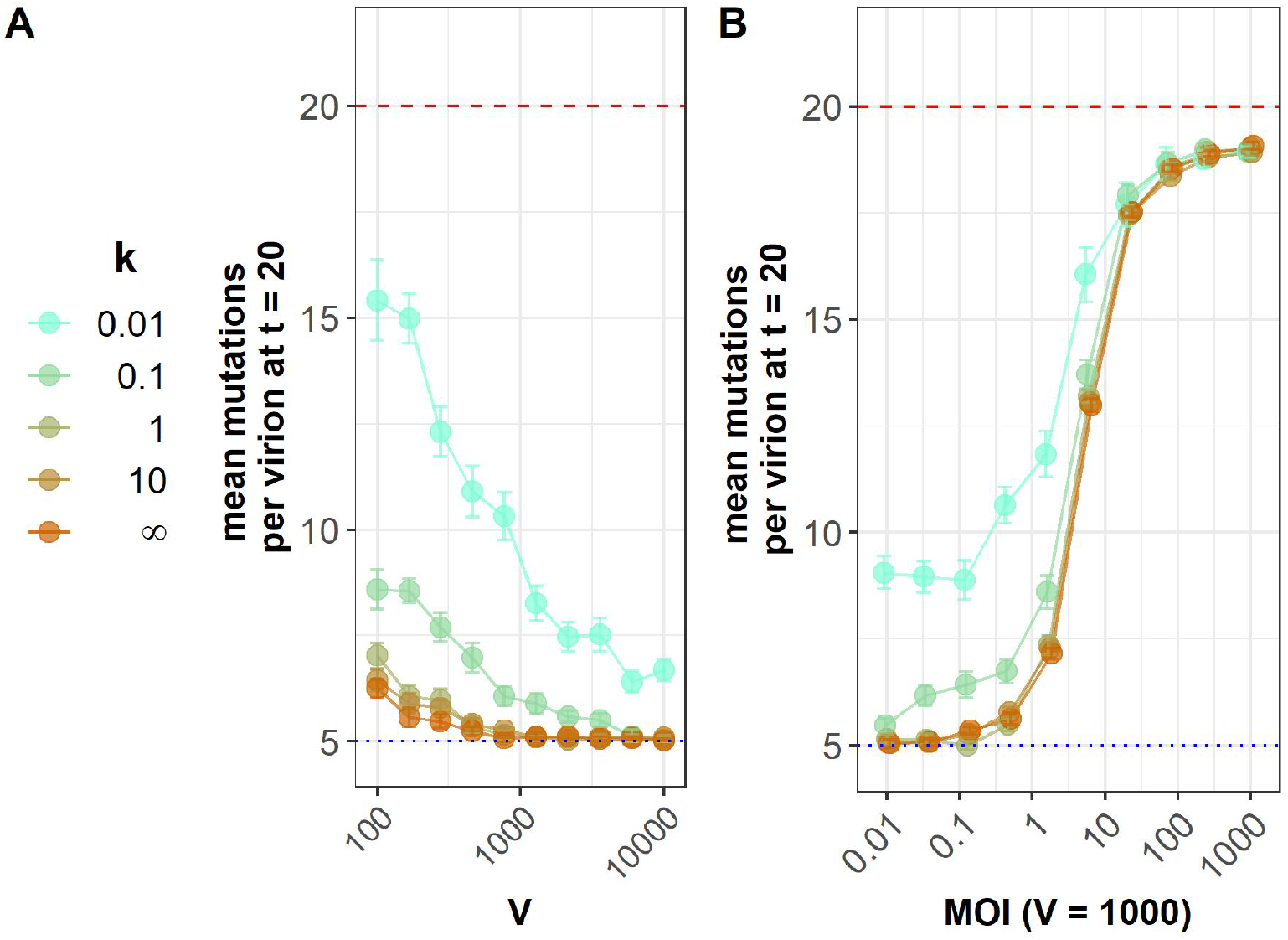
Stochastic heterogeneity increases deleterious mutation accumulation. All panels show mean number of deleterious mutations after *t* = 20 generations of within-host infection. Each data point shown is the average of 20 replicate simulations with error bars showing the standard deviation. Red dashed lines show the theoretical expected mutation accumulation at selective neutrality (*Ut*). Blue dashed lines show the average number of mutations for an infinite viral population size at its mutation-selection balance (*U/s*). Parameters are *U* = 1, *s* = 0.2, *y* = 1, with values of *k* indicated by color. **(A)** At a constant MOI of 0.1, stochastic heterogeneity increases mutation accumulation the most at intermediate population sizes where it increases the rate of Muller’s ratchet. **(B)** In these simulations, the virus population size is kept constant at *V* = 1000 and cell population sizes are modified to change MOI. Different *k* are tested.

### 2 Input-Dependent Heterogeneity

**Figure 2:**
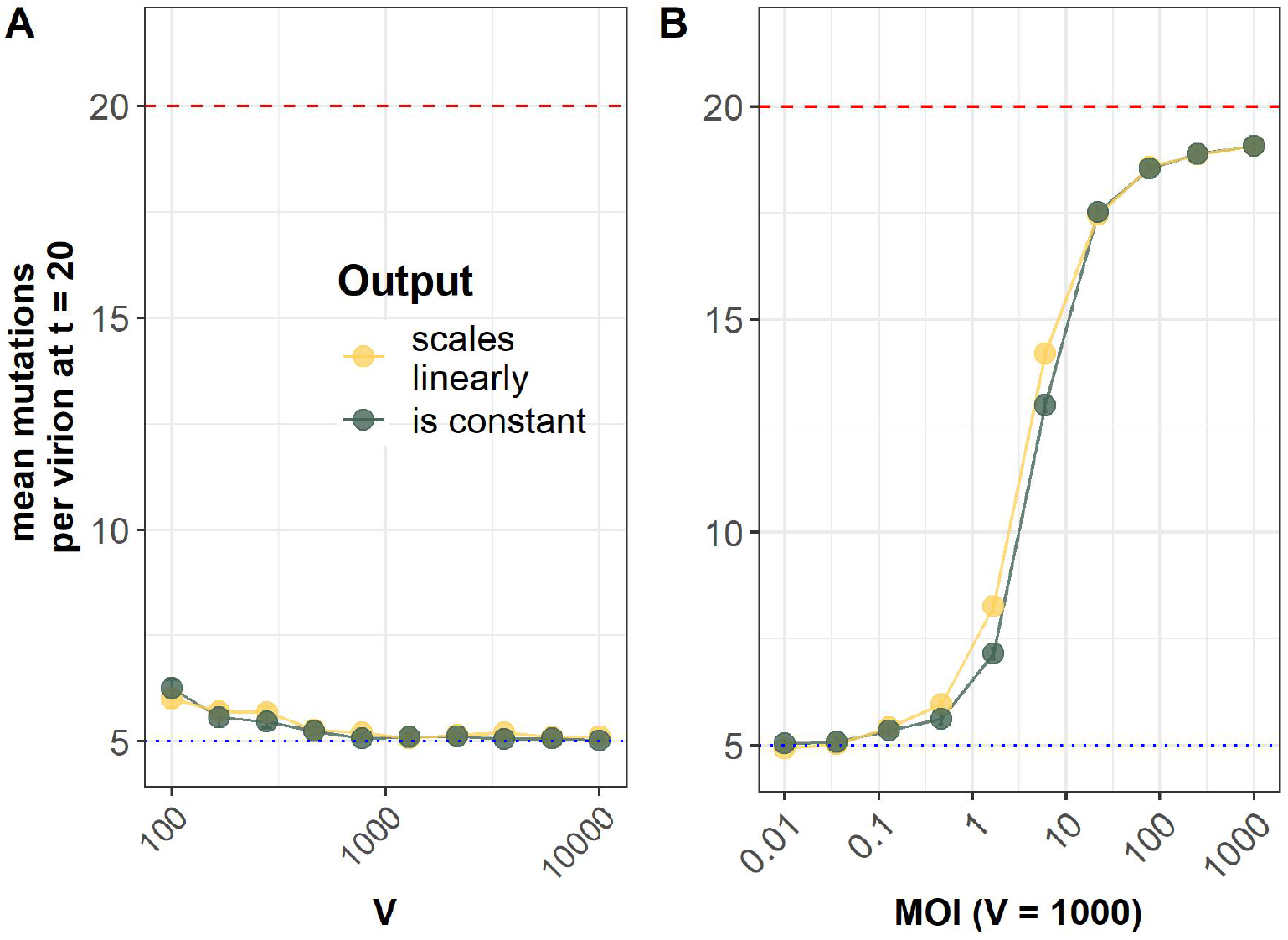
Input-dependent cellular fitness values affect mutation accumulation patterns. The black line represents simulations where there is no input-output relationship (i.e. the base model). The yellow line represent simulations where we have implemented our input-dependent model. We use the per genome mutation rate *U* = 1 and the fitness cost of mutations *s* = 0.2. Each data point shown is the average across 20 replicate simulations with error bars showing the standard error. Red dashed lines show the theoretical expectation of mutation accumulation at selective neutrality (*Ut*). Blue dashed lines show the expectation of mutation accumulation for an infinite viral population size at its mutation-selection balance (*U/s*). **(A)** Average number of mutations harbored by an individual virion at generation *t* = 20. With a constant MOI *V/C* = 0.1, different viral and cell population sizes are used to test the role drift. **(B)** Average number of mutations harbored by an individual virion at generation *t* = 20. The virus population size is kept constant at *V* = 1000 and cell population sizes are modified to change MOI.

